## Supplemental Figures for "Sexually dimorphic renal expression of *Klotho* is directed by a kidney-specific distal enhancer responsive to HNF1b"

### Figure and table legends

**Supplemental Fig. 1 No eRNAs at are detected at the *Klotho* enhancer locus.** ChIP-seq and RNA-seq were performed as described in the materials and methods section. Alignment of the RNA-seq to the PolII tracks indicates absence of enhancer RNA.

**Supplemental Fig. 2 Mouse weight is not impacted by enhancer 1 knockout.** Random litters of male, female, WT and E1 KO mice (n=8) at 3-4 months old were weighed. Unlike other Klotho knockout models, there is no significant difference in weight and mice survive into adultohood. Bar = SEM.

**Supplemental Fig. 3 There is no indication of female sex hormones impacting Klotho levels.** Renal Klotho mRNA was measured in 3-week old mice (A, n=4, data excluded due to undetectable *Klotho* expression) and in 8-week old mice ovariectomized at 3-weeks old (B, n=4). Sexual dimorphism of Klotho is detectable in prepubescent mice and ovariectomy does not impact mRNA levels. E2 deletion in females has no effect on Klotho expression (C, n=4.) Bar = SEM

**Supplemental Fig. 4 E1 deletion does not impact blood biochemistry.** Serum biochemistry panel was performed as described in materials and methods section. No differences were observed between any of the experimental groups. A-I – n=4. Bar = SEM

**Supplemental Fig. 5** Control loci for RNA-seq analysis; housekeeping genes expressed in the kidney, not directly related to Klotho and displaying similar expression levels across genotypes.

**Supplemental Fig. 6** Representative photographs of injured kidneys and contralateral controls 28 days after unilateral kidney ischemia reperfusion surgery in male and female WT and E1/E2 mice. Bar = 50μm.

**Supplemental Table 1** sgRNA and primer sequences used to construct and genotype E1, E2 and E1/E2 mice.

**Supplemental Table 2** Klotho E1 and E2 deletion sequences.

**Supplemental Spreadsheet 1** Gene counts and differentially expressed genes in WT and E1 KO male and female mice.

### Supplemental Figure 1

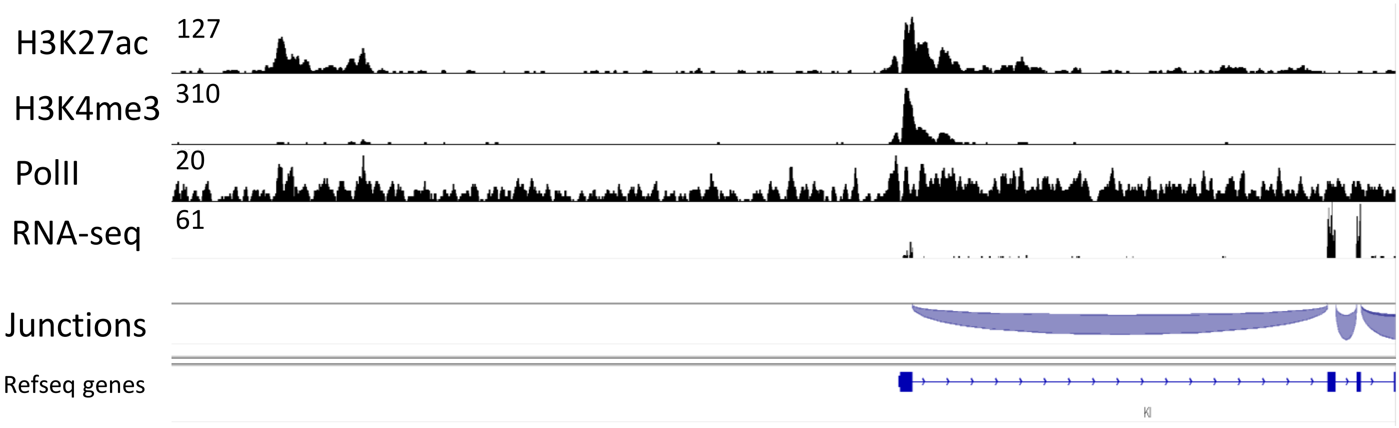

Supplemental Figure 2

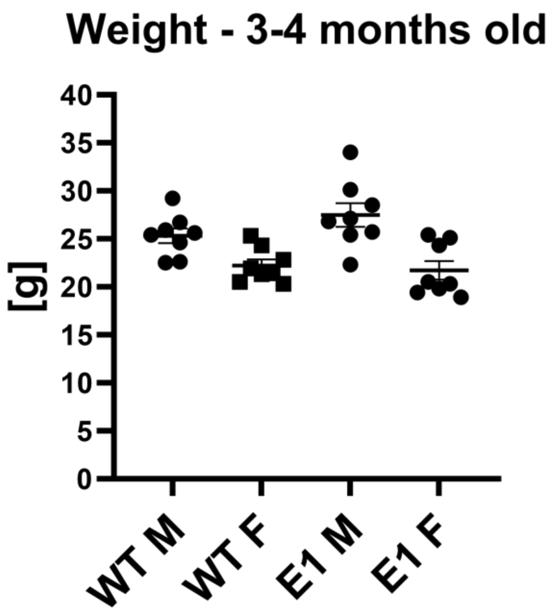

Supplemental Figure 3

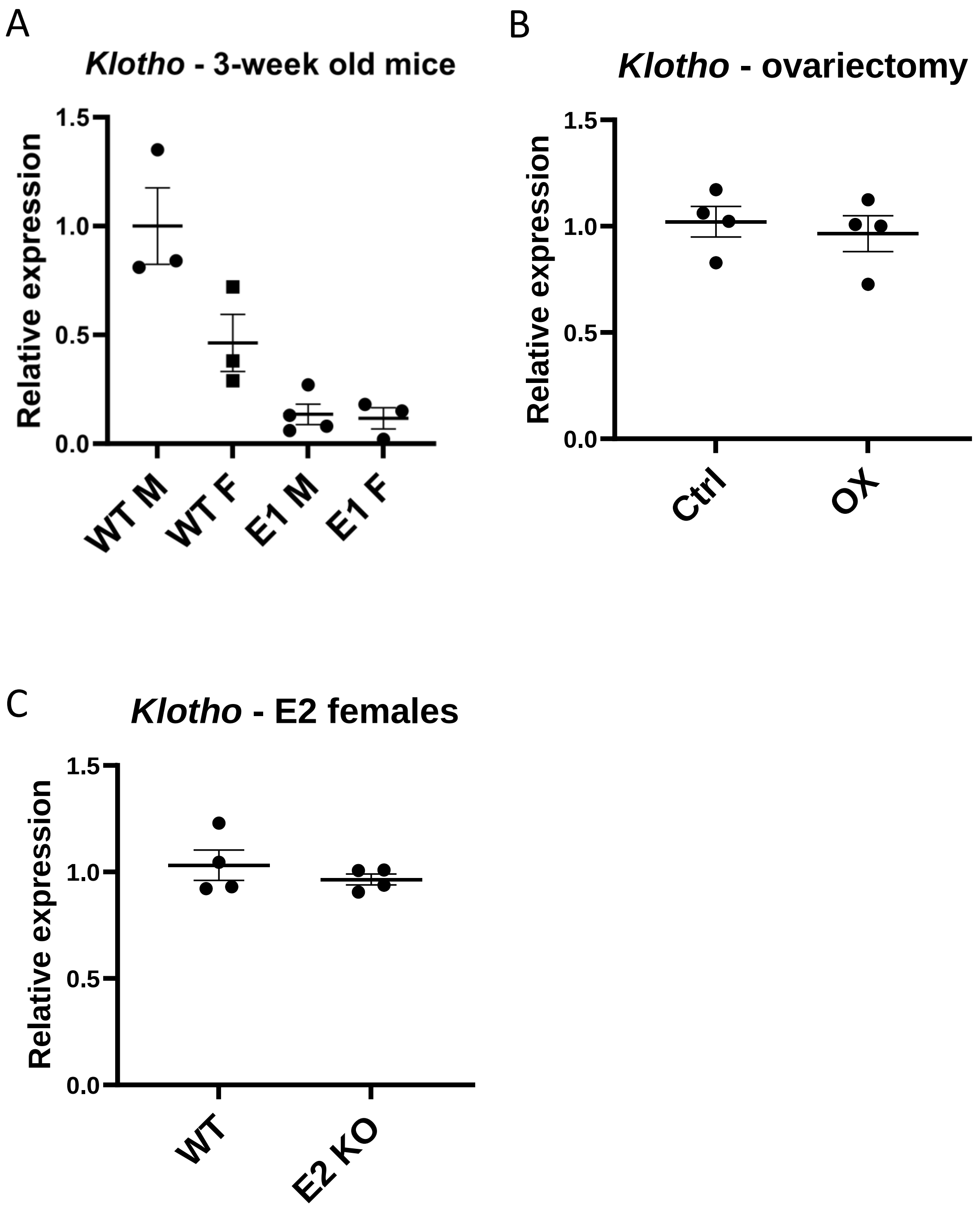

Supplemental Figure 4

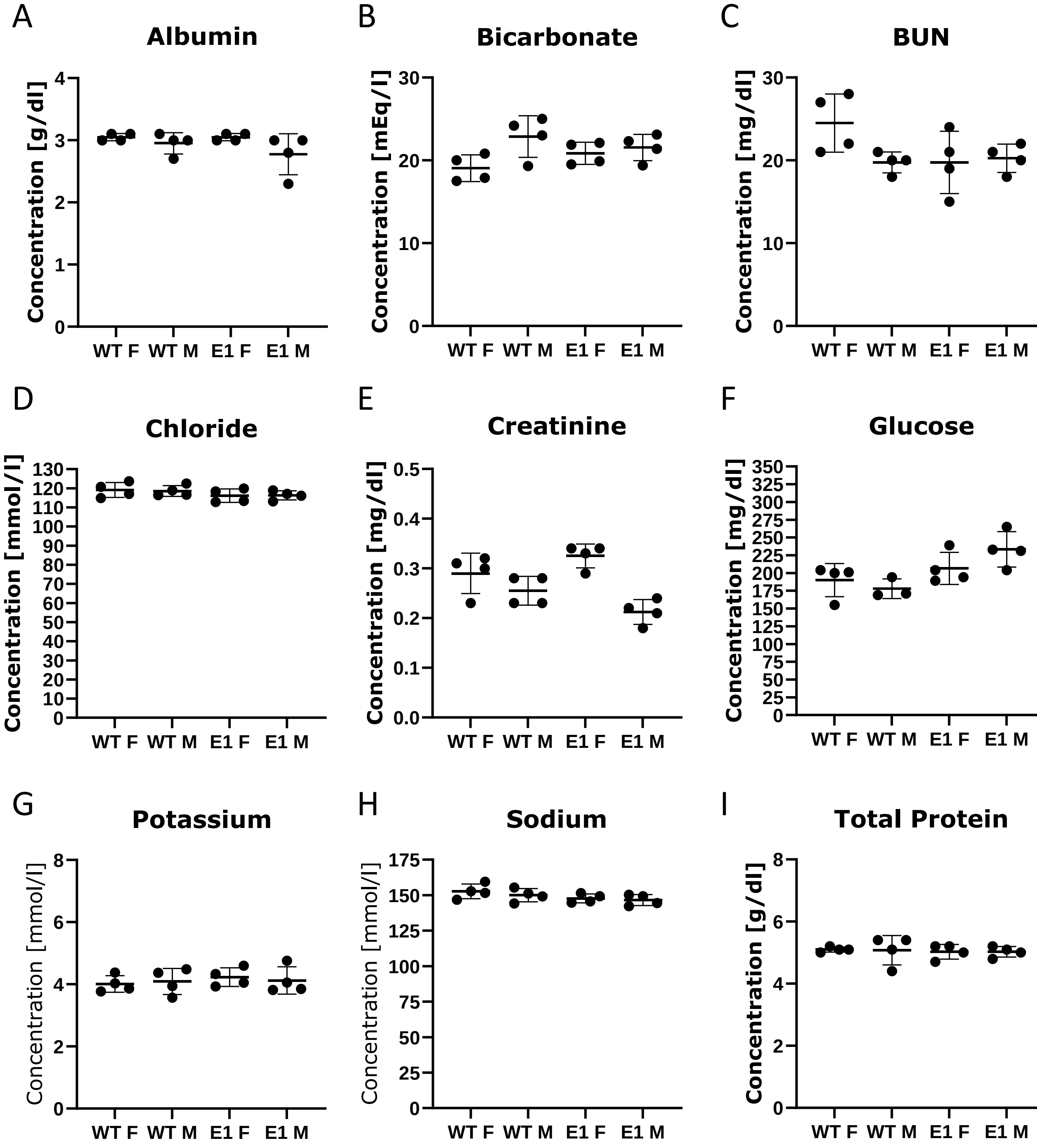

### Supplemental Figure 5

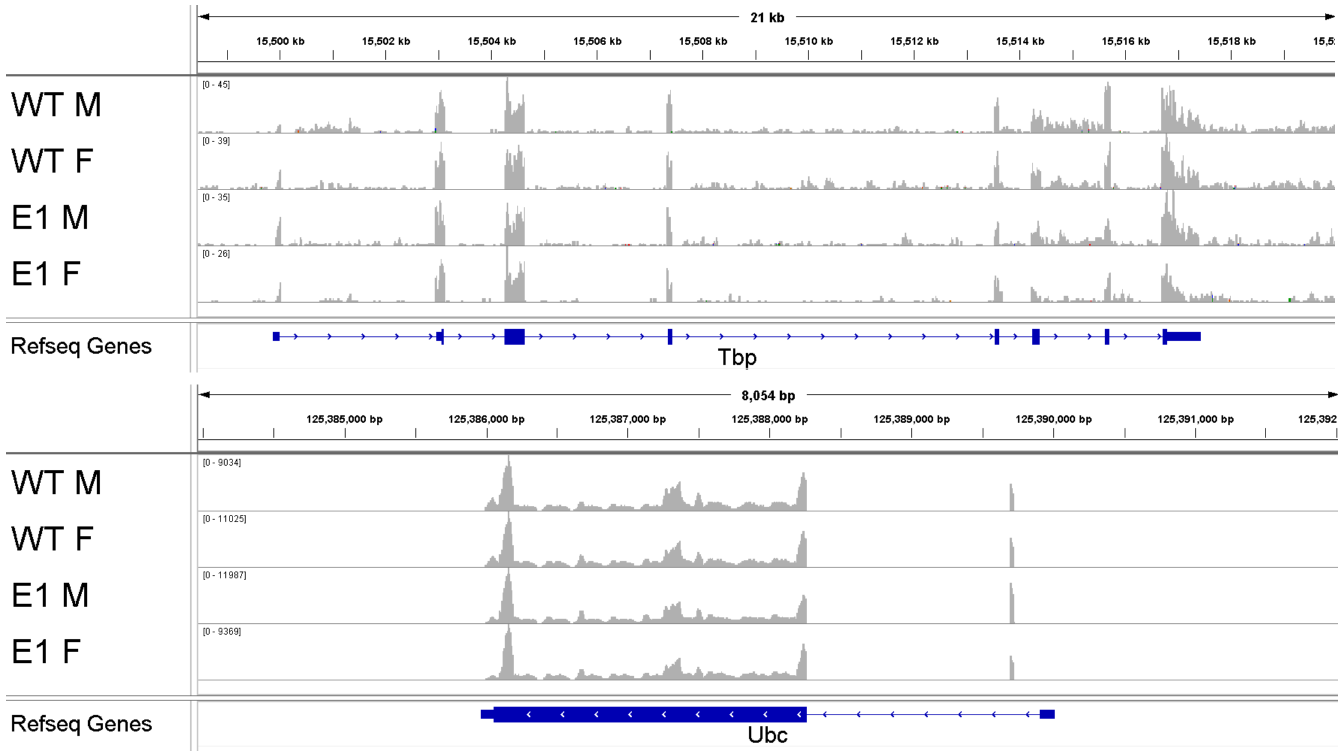

### Supplemental Figure 6

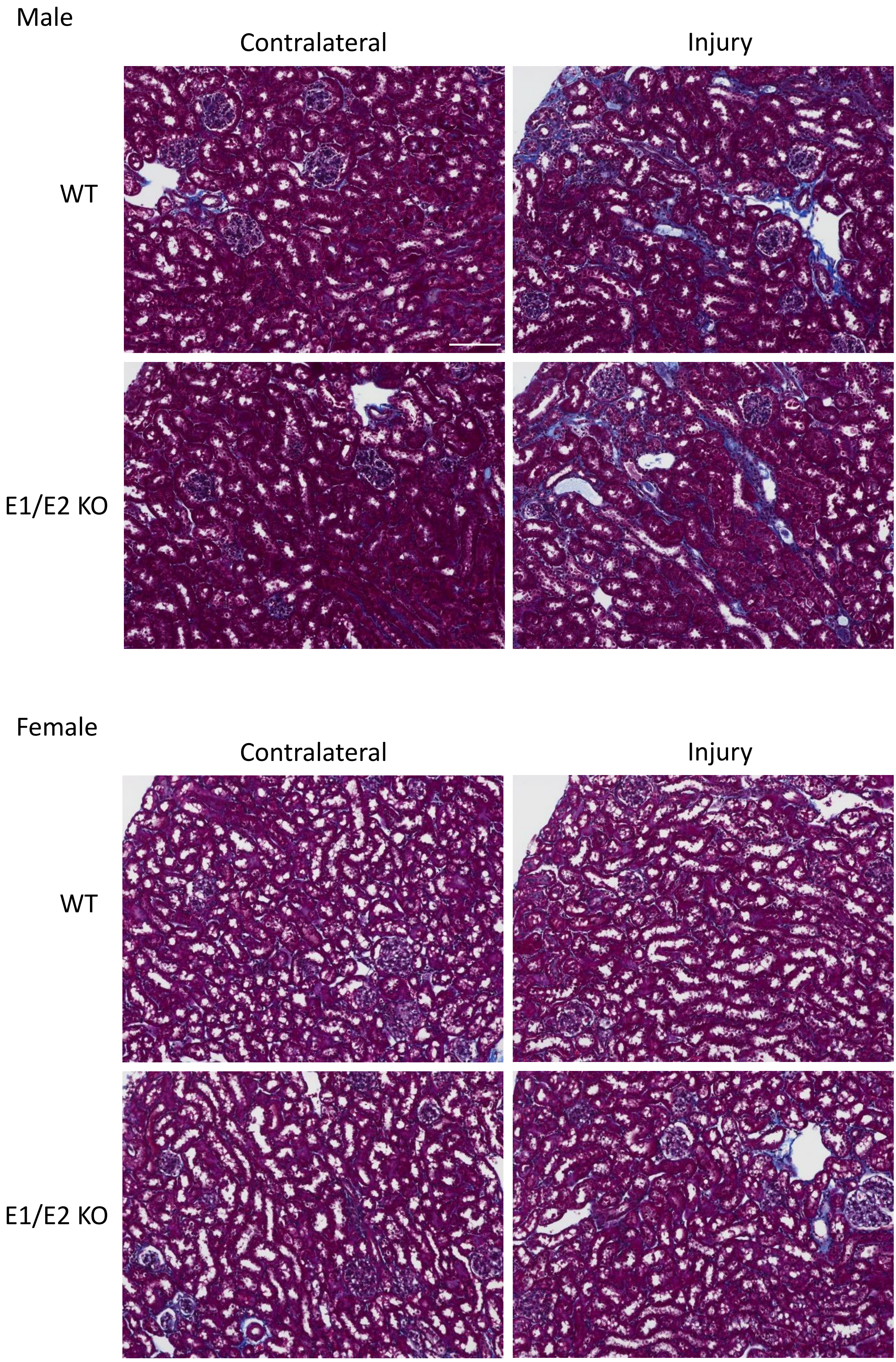

Supplemental Table 1

| | **Sequence** |
|---|---|
| E1 sgRNA | CTTCTTTCAGTGTGTCGCTTAAA |
| E2 sgRNA 1 | ACCCTGATGCACCTCTGAAG |
| E2 sgRNA 2 | TATCAGGTGGGATGAAGCCA |
| E1 Forward Primer 1 | GAACTCAGAAATCTGCCTGCTCC |
| E1 Reverse Primer 1 | GAGTAGCTGGGACTATGGTAATGC |
| E2 Forward Primer | CTATTCAGTGGTGTCGTTGGTCC |
| E2 Reverse Primer | TTGTTCTGTCCACCATCTCTTCCC |

### Supplemental Table 2

| Mouse strain | Deletion sequence |
|---|---|
| E1 KO (1744 bp del) | CTTAACTCATCTGTTGATGGACTGTTATCCTGGCTGTTGAGAACAAAGCTTCAATAAATATGGGAGTGAAACTGCCCTTTG ATGTACTGAGTTATTTTCCTTTGGGACTGGGAATTGCTGGATCACATGGAAATTTGTTCTTCATATTCTTGGTGACTGCACT GGTTTCCATAATGACAGGACTAATATTAAGTCATGGTTTGAAGACAGCAGTCATCTTAGAGGTGTGTGGCCCTAGCACCC TTCCCTTCCTCATGTGTAGCCCTAGGAAAGCAAACAAATAGTACAGGCTTCCCTATATTAGAAAGACCACAGCTGGGCTG GAGAGATGGCTCAGTGGTTAAGAGCATTAACTGCTCTTCCAAAGGTCCTAAGTTCAAATCCCAGCAACCACATGGTGGCT CACAACCATCTGTAATGAGATCTGATGCCCTCTTCTGGTGTGTCTGAAGACAGCTACAGTGTATTTACATATAATAAATAA ATCTTAAAAAAAAAAAAAAGAAAAAAAAAAAGACCATGGTTGTGTACTCTCTCACCTCCACAAGCCATGAGGATCAAAGGA GACAAACTGGGAAGACATGCTGGCAACTCAGTCACTCTGTGAATGTTTGCTGTTCTCTCAGATTGGACCTTACTCTGTATC TATCAAATTTGATATCTCCAGCAAGCCTAGTGAGTGTGGTTTGTCACTTAAGAACAGACTTAATTGGGGGGGGGGCTAAA TCTGGGAGTGTTAGTACTAAACATGTCTCTATGTGACTGTCTCACCAGGTCCAGGAATAATTACAAGTTTAGGGGTTGAG ATTTACTTGCATAGAGAGGGTAGGGATAGGGTGGGAAGGGTAAGAGGTTAGAAGGAAGGCAAAGGGTCACATGATGT CAAGTGCTGGAGAACACTGGGCAGTGTTCAGTGTGGATGGAGAGGAAGAAGTCAGGGGAATCACAGCACCCCATTCAC AGCTGAGCATCAGCTGGGTCCCGTGTCTGTGCTTGAGAGAGAGTCAGTAGTCAGTAGTCCTGGTGATAGGAGATGAGCT GCAGACAGAGGTGAGATACTCCCTGGCTGGCTCTCCTGCCTGAGTGCTCTCTGCACTTGCAGCCTCACGACTTTCCCTTGT ACTCTTGCACATGGCCATGGTAAAGGTTAACTGTGACCACCTGTCAAAGCCTGTTATATGTAGTTCCTCTCAGCCAGGCCT TATGTGAAGCCTGGGTGACTGCAGGAGGCAAAGCAGACATGGCCCCTGGACTCTTGGCACTCAGTGTCTGTGGGGGAG GTACACATGAAGACACAAGGAAATGCATACCCTAGAGCCAGGACACGTTGTGGAGGAGATGCCTGGAATCCTTGAGAGT TTAGAGACAGGTCTGCACACTGAAGGTGTGTGAAATAAAAATAAACAAACACACAAGAGCGGTACATTAAATTAAAAAA AAAATCTTGGCAAATACAAAGTGACTAGTTCTCACAGCAAGATCAAGTCTTGCTTTTAACTGAAGATGTTGGGGTTTTATT GTTTTAACAAATATGTGTGTTTTCCTCTTTCCGAGTTTGGAAAAACAATCAAGCAGACATCCCACAAAGTGCAGCCCTGGA AAACCTCGCCATCTGGACTGTCTGCTGGAAAAAACAGAAAGGTCACCCAGTGTGACTTGTCGAGCTTCACTAATTCCTGG AATTCCGTAGTCATTTCTCCAAATTAATTTTTAACTTCTTTCAGTGTGTCGCTTAAAATAAAATTT |
| E1 1001 bp del | GTGTGTCGCTTAAAATAAAATTTTTATCTCACTAAGGGGGGAAAAACAGTGGGAGGGGAAAAAGGATGGAGGGGTTGA CTTAACCTTTAAGTGTTAGCAGCACACAGAATTCAAAGTGCTTTACATAAATCAGAATCAGGGAAAAGCCAGGTGAGGTA GTGCATTACCATAGTCCCAGCTACTCAGGAGACTGAGCAGGAAAACCCACCCCAGTTTCAAGGCCAGCCTGCGCTACAGA GTGAGTTTAATGCTAGTCCAGGCTCTGCAGAGAGTTACTTTGTAACGGACTATAGAAATGATCCCTGAAAGTATAGTCTT GAGAGTTATTTGTATATTCCTCTTATCTTTTTTTTTTTTTTTTTTTTTTTTTTTTTTTGGTTTTTTTCGAGACAGGGTTTCTCT GTATAGCCCTGGCTGTCCTGGAACTCACTTTGTAGACCAGGCTGGCCTCGAACTCAGATTCACCTGCCTCTGCCTCCCGAG TGCTGGGATTAAAGGCGTGCACCACCACGCCCGGCCTTATTCCTCTTATCTTAGTATCTCCCAGAATTCCATGCCCCCTTAA CTACAGGTAAAAGTGAGACTTATCTACAGTAAAGCTACAGTGTGAAGCTCCTGGGCTGTCTTTGTCAACTTCTTCTCCACT CACATCATATTTGTTAATGAAACCTTTAGTGTTTATTCATTTATTTTATGTGTACAGATGCTTTTTGTTCTCTGGCATTTATAT ATGTGCACCATATCCATACAGTGTCCATTGGGGCCAGGCAAGGGTGTCAGATCCTATGAAACTGATTAAGGACAGTTTTG AGACATCCTGAGGGTGTTGGGAAGTGGACCCAGGTCCTCTGAAAGAGCAGCCAGTGCTTTTAACTGCTAAGCCATCTCTC ATGCCCAGCCATGGTCTTTCTAGTACAGGGAAGCCTGTGATAGAAGCACCTTCTATAGAAAGTGTCCAGTGAGTGTAGCT AGTGACTTCTAAGGTCAGTGGGTGTCTCA |
| E1 145 bp del | ATCTGGACTGTCTGCTGGAAAAAACAGAAAGGTCACCCAGTGTGACTTGTCGAGCTTCACTAATTCCTGGAATTCCGTAG TCATTTCTCCAAATTAATTTTTAACTTCTTTCAGTGTGTCGCTTAAAATAAAATTTTTATCTCAC |
| E1 31 bp del | ATTTTTAACTTCTTTCAGTGTGTCGCTTAAA |
| E2 KO | ACCCTGATGCACCTCTGAAGAGGAGGTCAGAGAGTTCACAGGTAAAAGACGAGGTTGGAAATACTTTTAAAGAATACAA GTTCACGGACCAGTGAGCCGCGGTTATGCATTTAGGAGCCGTCGCCTTTTGCGAGTGACAACTCACTGTCCACGTACTTG GGCAGGAGAGATAGAAGTTAATAAATGAACACGAAGGGACCCAGAAAAGAGCATTGCTGATAGAACACAGCAGAGCTG GGTGGATGCCAAACAGCCATGGGCACACCGCAGCGAAAGGAGGAACAACAGAAGCGCTTTCCAACAAATAACCAACAA ATTTGGTGCGGAGGGCAGAGAGACGACTGCTGAAGGGAGCTGATCGTTGTGCTGGGCCAGGAGGATTAAGATGAGTTT GCTGCTTATATCCAGACACTGTGCCATGTGCTCCCAGCTCGAGGGGGCTGAAGCCAGAACGCTGAGCAAGCCCAGCATG ACATGTGCACAGACGTGGTCCCTCTCCCTCTGCAAGAGAAGCCCCAATGCCTTCCTGGTTCCAAATCGCCTGAAATTGAG GTGTATGAAACTGGGTACTGGCTGGTAAATTTACCTTAGTAAGGCACGCTCTGAACTCCACTGCCGGTCAAAGGTGGATT AATTCAGATGACCTGGAGGGAAAGTTCACACGAATCACCTCTCTCTCTCTCTCTCTCTCTCTCTCTCTCTCTCTCTCGCTC TCGCTCTCGCTCTCTCGCTCTCTCTTGCTTTCTCTCTCTTGCTCTGTCTCTGTCTCTCTCTCTCTCTCGCTCTCTCGTTCTCTC GCTCTCTCGCTCTCTCGCTCTCTCGCTCTCTCGCTCTCTCGCTCTCTCGCTCTCTCGCTCTCGCTCTCGTTCTCATCTCATCCC ATGGCAGATGGGCGCACTCTGTTGTGGCTACAAAACTCAGGAACGCCCAGGCTTGCTAATTCAGGTATCAGGTGGGATG AAGCCAAGGG |
| E1/E2 KO (E2) | CGAGGTTGGAAATACTTTTAAAGAATACAAGTTCACGGACCAGTGAGCCGCGGTTATGCATTTAGGAGCCGTCGCCTTTT GCGAGTGACAACTCACTGTCCACGTACTTGGGCAGGAGAGATAGAAGTTAATAAATGAACACGAAGGGACCCAGAAAA GAGCATTGCTGATAGAACACAGCAGAGCTGGGTGGATGCCAAACAGCCATGGGCACACCGCAGCGAAAGGAGGAACAA CAGAAGCGCTTTCCAACAAATAACCAACAAATTTGGTGCGGAGGGCAGAGAGACGACTGCTGAAGGGAGCTGATCGTTG TGCTGGGCCAGGAGGATTAAGATGAGTTTGCTGCTTATATCCAGACACTGTGCCATGTGCTCCCAGCTCGAGGGGGCTG AAGCCAGAACGCTGAGCAAGCCCAGCATGACATGTGCACAGACGTGGTCCCTCTCCCTCTGCAAGAGAAGCCCCAATGC CTTCCTGGTTCCAAATCGCCTGAAATTGAGGTGTATGAAACTGGGTACTGGCTGGTAAATTTACCTTAGTAAGGCACGCTC TGAACTCCACTGCCGGTCAAAGGTGGATTAATTCAGATGACCTGGAGGGAAAGTTCACACGAATCACCTCTCTCTCTCTCT CTCTCTCTCTCTCTCTCTCTCTCTCTCGCTCTCGCTCTCGCTCTCTCGCTCTCTCTTGCTTTCTCTCTCTTGCTCTGTCTCTGTCT CTCTCTCTCTCTCTCGCTCTCTCGTTCTCTCGCTCTCTCGCTCTCTCGCTCTCTCGCTCTCTCGCTCTCTCGCTCTCTCGCTCTC TCGCTCTCGCTCTCGTTCTCATCTCATCCCATGGCAGATGGGCGCACTCTGTTGTGGCT |
